# Rapid non-destructive method to phenotype stomatal traits

**DOI:** 10.1101/2022.06.28.497692

**Authors:** Phetdalaphone Pathoumthong, Zhen Zhang, Stuart J. Roy, Abdeljalil El Habti

## Abstract

**Background:** Stomata are tiny pores on the leaf surface that are central to gas exchange. Stomatal number, size and aperture are key determinants of plant transpiration and photosynthesis, and variation in these traits can affect plant growth and productivity. Current methods to screen for stomatal phenotypes are tedious and not high throughput. This impedes research on stomatal biology and hinders efforts to develop resilient crops with optimised stomatal patterning. We have developed a rapid non-destructive method to phenotype stomatal traits in four species: wheat, rice, tomato and Arabidopsis.

**Results:** The method consists of two steps. The first is the non-destructive capture of images of the leaf surface from plants in their growing environment using a handheld microscope; a process which only takes a few seconds compared to minutes for other methods. The second is to analyse stomatal features using a machine learning model that automatically detects, counts and measures stomatal number, size and aperture. The accuracy of the machine learning model in detecting stomata ranged from 76% to 99%, depending on the species, with a high correlation between measures of number, size and aperture between measurements using the machine learning models and by measuring them manually. The rapid method was applied to quickly identify contrasting stomatal phenotypes.

**Conclusions:** We developed a method that combines rapid non-destructive imaging of leaf surfaces with automated image analysis. The method provides accurate data on stomatal features while significantly reducing time for data acquisition and analysis. It can be readily used to phenotype stomata in large populations in the field and in controlled environments.

## Background

Stomata are pores on the surface of leaves, stem and floral tissues of plants (1). Stomata play an essential role in plant growth and plant response to abiotic and biotic stress. Approximately 98% of carbon dioxide (CO_2_) uptake and water loss from the plant occurs through stomatal apertures (2). Stomata are dynamically regulated by environmental factors such as drought, heat, salinity, light, to name a few. When water supply is ample, stomata open to allow carbon dioxide (CO_2_) entry into the leaf for photosynthesis. Simultaneously, water is released to the atmosphere via transpiration (2, 3). When water supply is limited, plants close stomata to prevent water loss, which also results in reduced CO_2_ assimilation and subsequently growth. The balance between CO_2_ assimilation and water loss is primarily important for adaptation to environmental cues without compromising growth (2, 3). Stomata are also a gateway for pathogen entry into leaves and stomatal defence is a major physical defence mechanism in plant immunity (4, 5).

Stomatal gas exchange and its regulation are determined by stomatal morphology, density and sensitivity to the environment (6). Plants show a range of stomatal sizes and shapes on the leaf epidermis depending on the plant species and variety (7). Diversity in stomatal features allowed plants to adapt to a wide range of environments (8, 9). Considering a changing climate, the flexible and dynamic nature of stomatal traits makes them primary targets for improving crop productivity and stability. Despite extensive research on stomatal biology, current knowledge is poorly translated into the context of field experiments and outputs for breeders. Conventional methods to phenotype stomata are time-consuming and costly, and do not allow screening large populations for beneficial stomatal traits. A commonly used method to investigate stomatal traits is a nail polish method (NP method) (10-15). It consists of applying nail polish on the leaf surface to get a leaf imprint, waiting for the polish to dry, carefully pealing off the polish, examining the leaf imprint under a light microscope, taking and analysing images to determine stomatal traits. In addition to the length of time it takes to obtain a nail polish imprint, a common issue with this method is the unavoidable presence of air bubbles that interfere with stomata imaging (16), which can result in missing data and repeating the imprinting stage. Manual measurement of stomatal traits is time-consuming and inevitably introduces inaccuracies. Altogether, obtaining images of stomata from leaf imprints and analysing data is time-consuming and can take up to 40 min per sample, which limits the number of samples that can be analysed at any one time, making it impractical for phenotyping large mapping populations or diversity panels.

Recently, a number of methods have been published which address the issues with manual processing by including a machine learning algorithms that automatically detect and analyse stomatal features (16-20). While this improvement significantly accelerated image analysis and offers an effective substitute to manual analysis, these methods are limited by providing either information on stomatal number, size or aperture in one species in isolation; having data on all stomatal traits is desirable to have a comprehensive view on stomatal phenotype. In addition these faster image processing methods still depend on traditional time consuming approaches to obtain images, such as nail polish methods that require indirect leaf imaging from leaf imprints. Use of nail polish methods are impractical when working with large populations and having to take thousands of measurements from greenhouse or field grown plants, which require higher throughput image acquisition and analysis. To overcome indirect leaf imaging, approaches of directly taking images of leaves by bringing harvested plant material or whole plants to a microscope have been trialled (21-23). Direct leaf imaging further accelerates stomatal phenotyping and demonstrates the potential of high-throughput stomata phenotyping tools in revealing novel insights on the genetic basis of stomata-related traits (24). However, these methods are typically destructive or only provide partial information on stomata number and/or aperture.

Facing the need for a high-throughput method to phenotype stomatal traits in large populations and identify favourable stomatal traits, we developed a rapid non-destructive method for phenotyping stomata at large scale by combining a portable handheld microscope (HHM) for direct imaging of a leaf surface, with a machine learning model for automated stomata analysis (Fig. 1). This method was tested on four species: wheat, rice, tomato and Arabidopsis.

**Figure 1.**
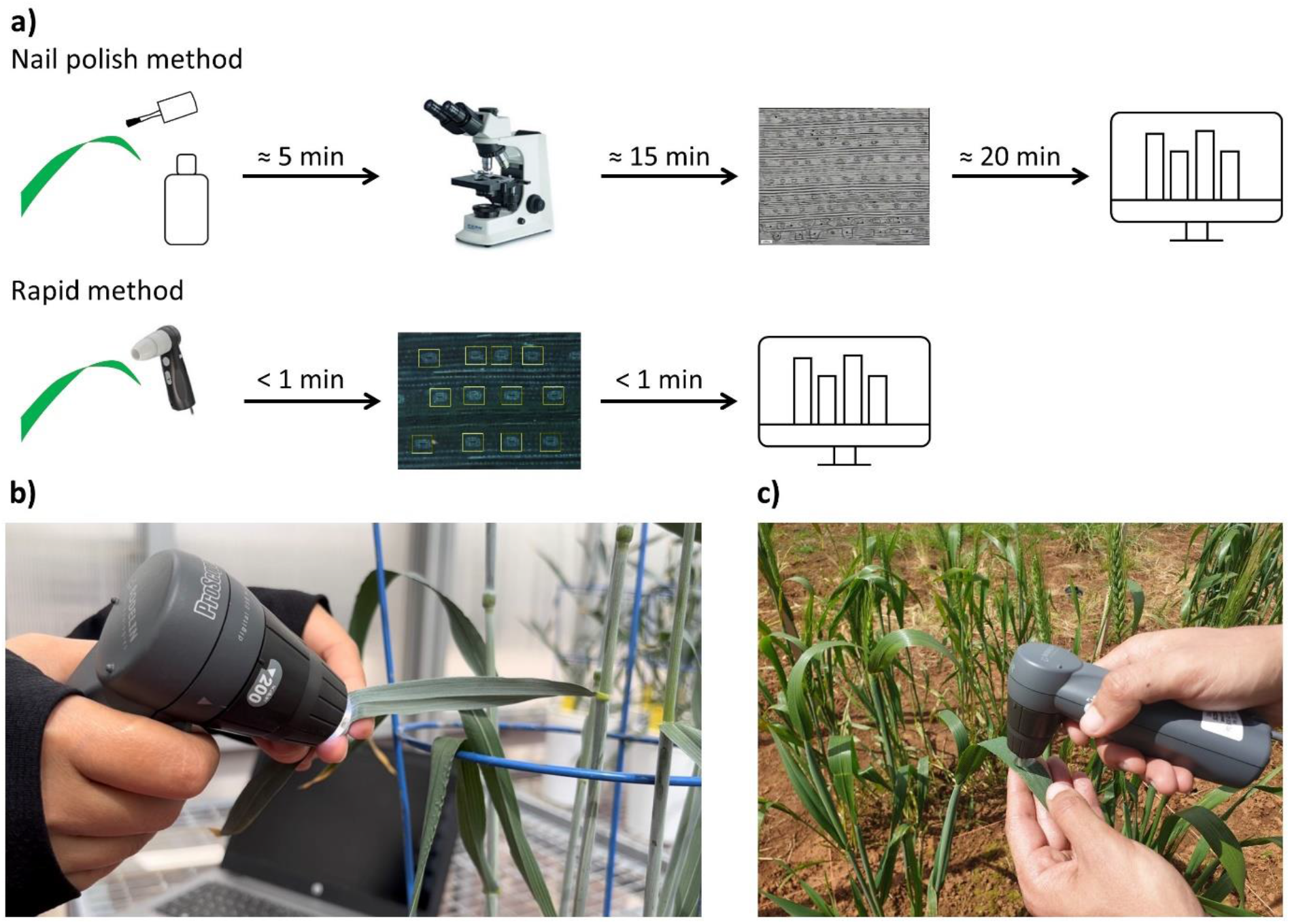
Overview of the rapid stomata phenotyping method (a) and experimental setup in controlled environment (b) and in the field (c).

## Materials and Methods

### Plant material

Four plant species were tested in this work: two monocotyledons (wheat - *Triticum. aestivum* cv. Cadenza and Gladius, and rice - *Oryza sativa* cv. R12) and two dicotyledons (tomato - *Solanum lycopersicum* cv. Sweetbite and Mighty Red, and Arabidopsis - *Arabidopsis thaliana* cv. Columbia). Wheat and tomato plants were grown in 20 cm pots containing UC Davis soil mix (50% peat and 50% sand) in a glasshouse located at the Waite campus (South Australia, 34°58’16.72”S latitude 138°38’23.17”E longitude) under natural photoperiod and 22°C/15°C day/night. Wheat was grown from June to October 2021 and tomato plants from July to November 2021. Rice was grown from September to December 2021 at 29°C/21°C day/night in 15 cm pots containing UC David soil mix. Arabidopsis was grown in a growth cabinet at 23°C/19°C day/night under a 12h photoperiod with a light intensity of 200 μmol m^−2^ s^−1^. Arabidopsis plants were grown in 8 cm pots containing Arabidopsis mix soil (Coir 3.6L, Perlite 3.6L & Sand 0.25L).

### Stomata imaging using the handheld microscope method

A handheld microscope (ProScope HR5, Bodelin, USA) was used to directly take images of plant leaves which were still attached to the plant. Four magnifying objectives were tested: 100× lens (field of view: 2.87 × 2.17 mm; resolving power: 4 microns; pixel density: 1mm = 198 pixels), 200× (field of view: 1.36 × 1.03mm; resolving power: 2 microns; pixel density: 1mm = 415 pixels) and 400× (field of view: 0.75 × 0.57 mm; resolving power: 1 micron; pixel density: 1mm = 652 pixels). Images were captured using ProScope Capture v6.14 software. Arabidopsis images taken with a HHM were cropped into 0.3 × 0.3 mm images to exclude areas which were out of the focal range.

### Stomata imaging using a nail polish method

For comparison with the HHM method, leaf imprints were collected by applying nail polish on the adaxial and abaxial leaf surface of 4-month-old wheat, 2-month-old rice, 3-month-old tomato and 2-month-old Arabidopsis plants. Nail polish (Insta-Dri Top Coat, Sally Hansen, USA) was applied on the leaf surface in a thin layer, and left to air dry for 5 minutes. Dry nail polish was peeled off using clear tape (Crystal clear office tapes, Winc, Australia) and was placed on a microscope slide. Stomata on leaf imprints were observed using a light microscope (Nikon Ni-E compound microscope, Tokyo, Japan) connected to a DS-Ri1 colour cooler digital camera and a 40× objective. Images of leaf imprints were taken using NIS-elements software (Nikon).

### Machine learning model training Stomata detection model

Images taken with the HHM were used to train the machine learning model to detect stomata for each species and magnification used. Open-source software LabelImg (25) was used to annotate images by labelling a bounding box around individual stomata. The annotated images were uploaded to roboflow platform (26) for image augmentation. YOLOv5 algorithm (27) was used to train the model for stomatal detection. Accuracy of stomata detection models was assessed with YOLOv5 using three parameters: precision, recall and F1 score. Precision is defined as

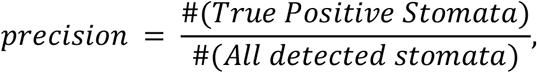

Recall is defined as

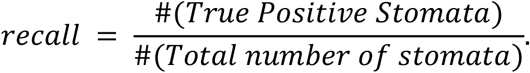

F1 score is the harmonic mean of precision and recall and is defined as

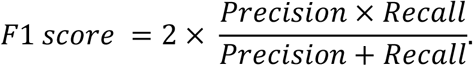

### Stomata measurement model in wheat

Data from the stomata detection model were used to extract individual stomata from the HHM images. Instance segmentation was done using Roboflow platform by manually outlining stomatal perimeter and aperture of individual stomata. Detectron2 platform (28) was used to train the model for measurement of stomatal area and aperture. Data on stomatal traits measured using the developed machine learning models were compared with data measured manually on 100 images. The number of stomata was counted visually. Stomatal size and aperture were measured manually using Fiji (28) by outlining stomatal perimeter and aperture.

## Results

### Optimal magnification for each species

To assess the suitability of each magnification in providing accurate data on stomatal traits, three magnifying objectives were tested: 100×, 200× and 400×. The suitability of a magnification in quantifying stomatal traits depended on the size of stomata in each species (Fig. 2). In wheat, the large field of view of the 100× magnification allowed the counting of stomata over a large area, but the lower resolution did not allow accurate measurement of stomata size (Fig. 2a). The intermediate specifications of 200× magnification enabled both counting the number of stomata in a smaller area on the wheat leaf and could be used to determine stomatal size (Fig. 2b). The high resolution of 400× magnification was optimal for determining stomatal size and aperture in wheat (Fig. 2c). Stomata in rice, tomato and Arabidopsis are significantly smaller than stomata in wheat and therefore only the 400× magnification could be used to determine the number and size of stomata (Fig. 2 d-f).

**Figure 2.**
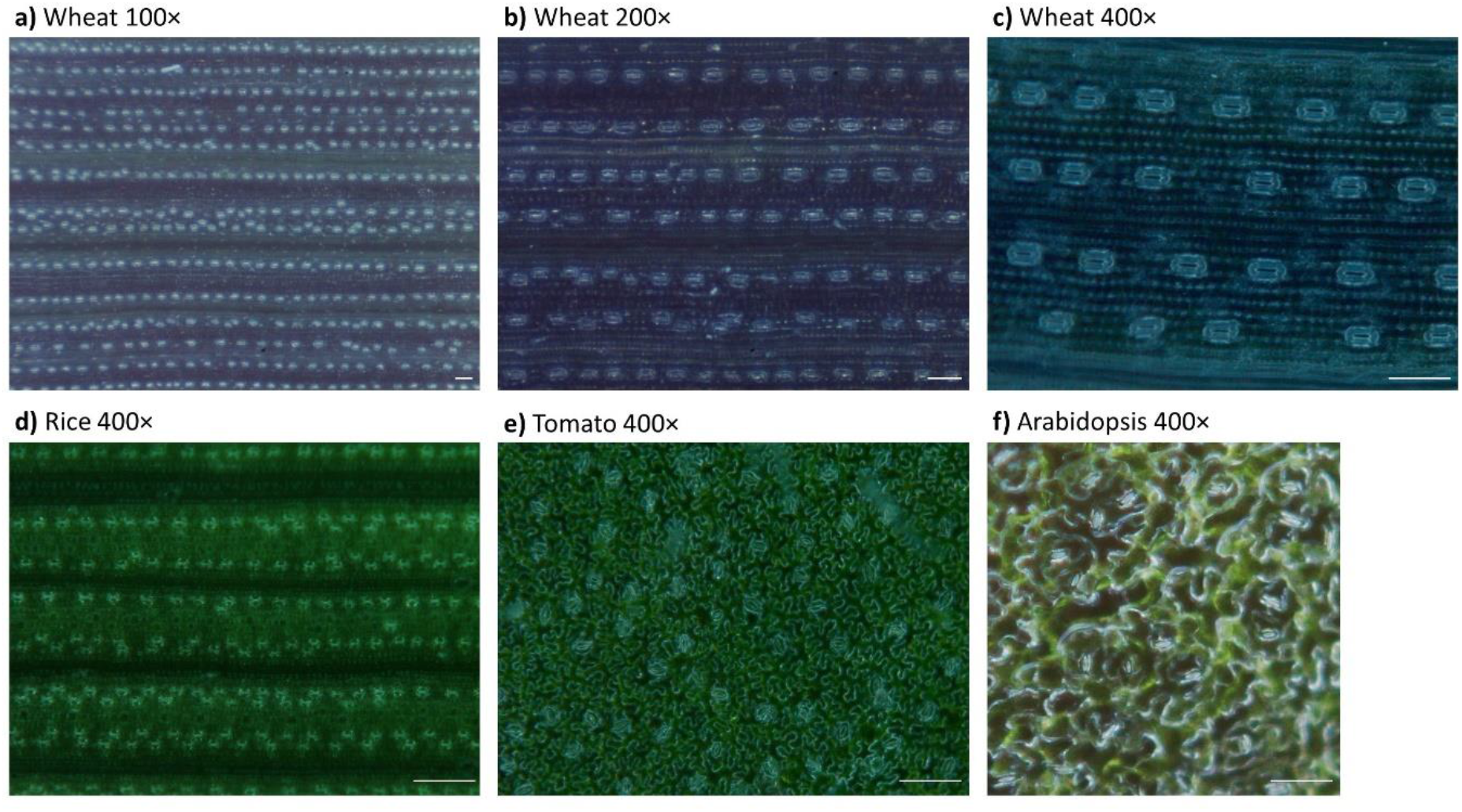
Handheld microscope images of wheat leaves with 100× (a), 200× (b) and 400× (c) magnification, rice (d), tomato (e) and Arabidopsis (f) leaves with 400× magnification. Scale bar = 100 μm.

### Comparison between nail polish and handheld microscope images

The HHM provided good quality images faster than the nail polish method. Although leaf imprint images taken with a light microscope were generally at higher resolution than the HHM images, air bubbles were frequently present in imprint samples (Fig. 3). Handheld microscope images of wheat leaves using 400× magnification were at high resolution given the large size of stomata in this species and allowed automated measurement of number of stomata, stomatal size and aperture (Fig. 3a). In rice, tomato and Arabidopsis leaves, images taken with the HHM were at a lower resolution compared to light microscope images given the smaller size of stomata in these species, but stomata were distinct enough to allow for automated detection of stomata and measurement of stomata number and size (Fig. 3b-d). Rice leaf imprints were uneven and required taking two images at different focus to visualise all stomata present on the leaf. The HHM allowed the observation of all rice stomata in one image (Fig. 3b). Given their smaller size, stomata in rice leaves could only be observed using the 400× magnification but image quality was still suitable for machine learning. In tomato, leaf imprint images were as clear as images taken with the HHM and stomata could be observed with a 400× magnification (Fig. 3c). In Arabidopsis, leaf imprint images were clearer than HHM images, but stomata were distinct enough to be detected and measured by the machine learning model (Fig. 3d). Arabidopsis presented a unique problem in that cells towards the periphery of the image were out of focus – an issue not encountered with wheat, rice or tomato. Due to this HHM images of Arabidopsis leaves were trimmed to include only the areas in focus before image processing took place.

**Figure 3.**
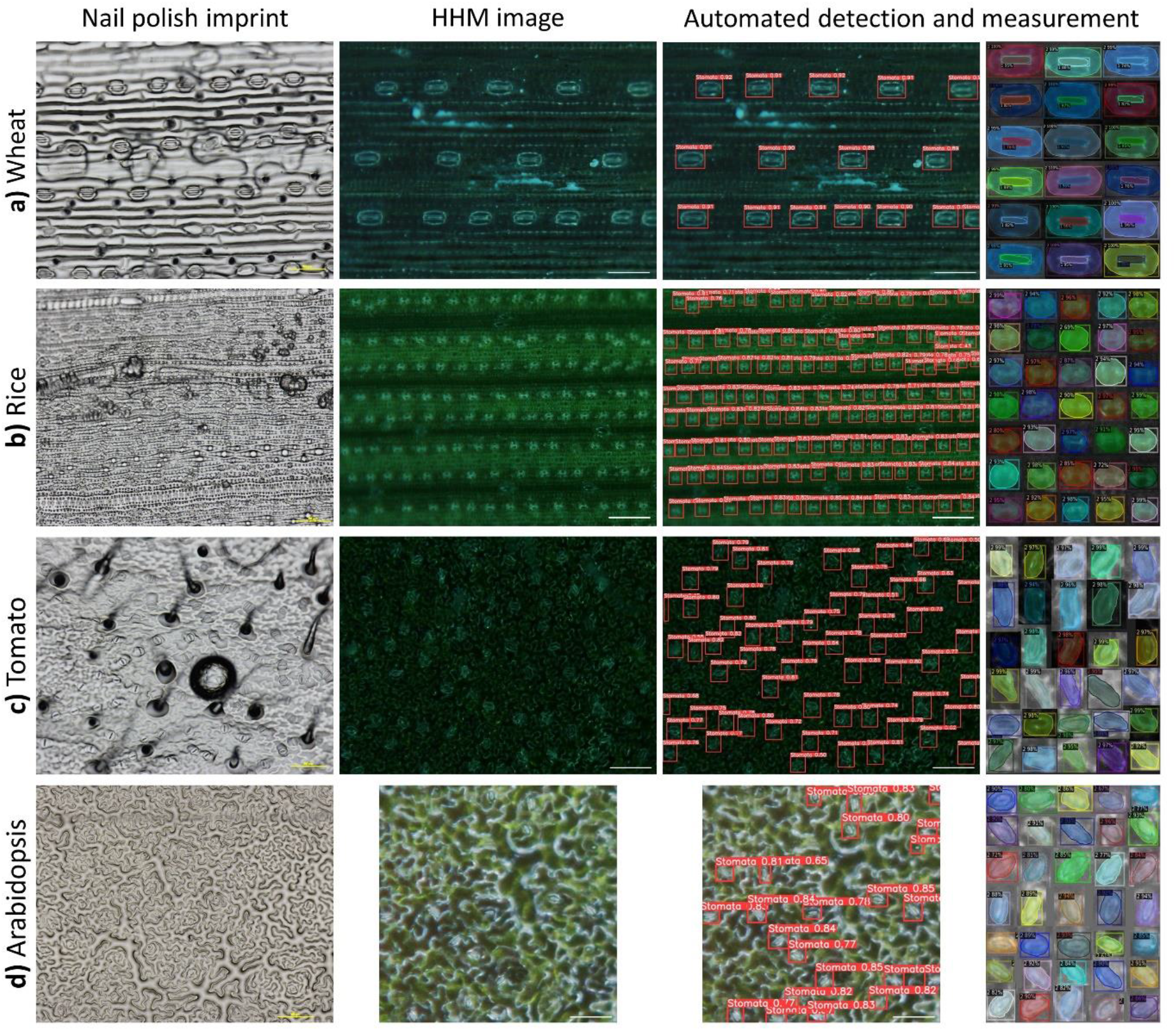
Qualitative comparison between images from nail polish imprint images and handheld microscope images of wheat (a), rice (b), tomato (c) and Arabidopsis leaves (d), and automated detection and annotation of stomata and aperture. Scale bar = 100 μm.

### Data accuracy of detection and measurement models

The machine learning models were highly accurate in detecting and measuring stomata. Model accuracy depended on the species and the magnification used (Table 1, Supp. Fig. 1). In wheat, model accuracy was highest with 400× images (F1 = 0.99) compared to 200× (F1 = 0.91) and 100× (F1 = 0.89), as stomata are more distinct at higher magnification. When comparing across all species using the 400× magnification, stomatal detection in wheat images had the highest accuracy compared to rice, tomato and Arabidopsis (F1 = 0.99, 0.90, 0.87 and 0.76, respectively).

**Table 1.** Detection models statistics.

| Species | Magnification | Precision | Recall | F1 score |
| --- | --- | --- | --- | --- |
| Wheat | 400× | 0.993 | 0.998 | 0.995 |
| Wheat | 200× | 0.898 | 0.917 | 0.907 |
| Wheat | 100× | 0.977 | 0.815 | 0.889 |
| Rice | 400× | 0.868 | 0.926 | 0.896 |
| Tomato | 400× | 0.836 | 0.913 | 0.873 |
| Arabidopsis | 400× | 0.775 | 0.742 | 0.758 |

Stomata detection and measurements using the machine learning models were highly accurate, meaning that the developed models were able to detect nearly all of the stomata in an image and accurately measure stomatal size and aperture (Fig. 4). There was a strong correlation between measurements obtained by the machine learning models and those obtained manually. In wheat, data from machine learning models were highly correlated with manual measurements for stomata number (R^2^ = 0.99), stomatal size (R^2^ = 0.87) and stomatal aperture (R^2^ = 0.94) (Fig. 4a-c). In rice, data from machine learning models were highly correlated with manual measurements for stomata number (R^2^ = 0.99) and stomatal size (R^2^ = 0.74) (Fig. 4b). In tomato, data from machine learning models were highly correlated with manual measurements for stomata number (R^2^ = 0.99) and stomatal size (R^2^ = 0.88) (Fig. 4c). In Arabidopsis, data from machine learning models were highly correlated with manual measurements for stomata number (R^2^ = 0.98) and stomatal size (R^2^ = 0.75) (Fig. 4d). The accuracy of detection and measurement model in wheat, rice, tomato and Arabidopsis allowed rapid identification of contrasting stomatal number and size; high stomatal density with smaller stomata, and low stomatal density with larger stomata (Fig. 5).

**Figure 4.**
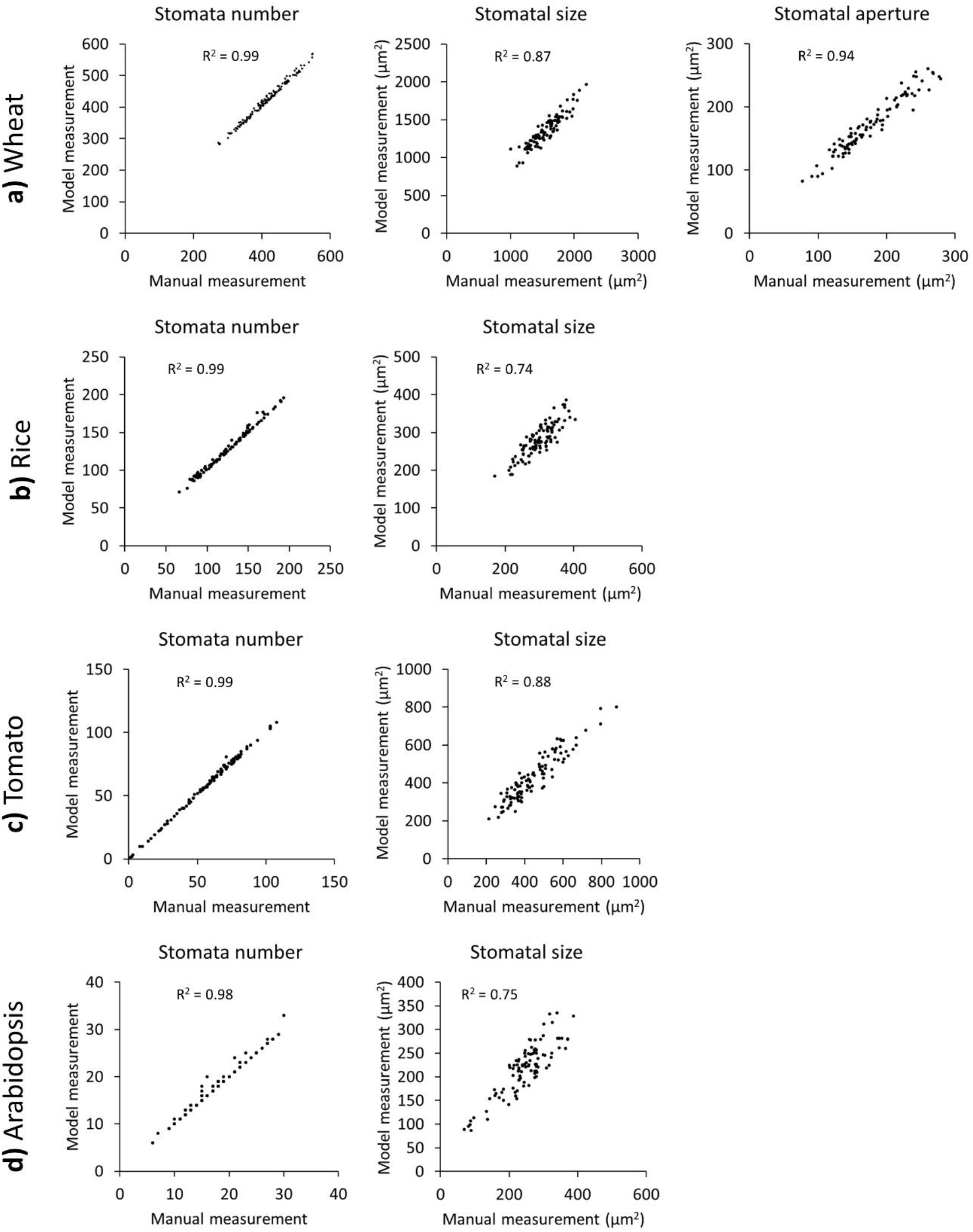
Scatterplots of stomatal traits comparing data measured by machine learning models with manual measurements of 100 images of wheat (a), rice (b), tomato (c) and Arabidopsis leaves (d). R^2^ is the coefficient of determination of a linear regression between computed and manual counts.

**Figure 5.**
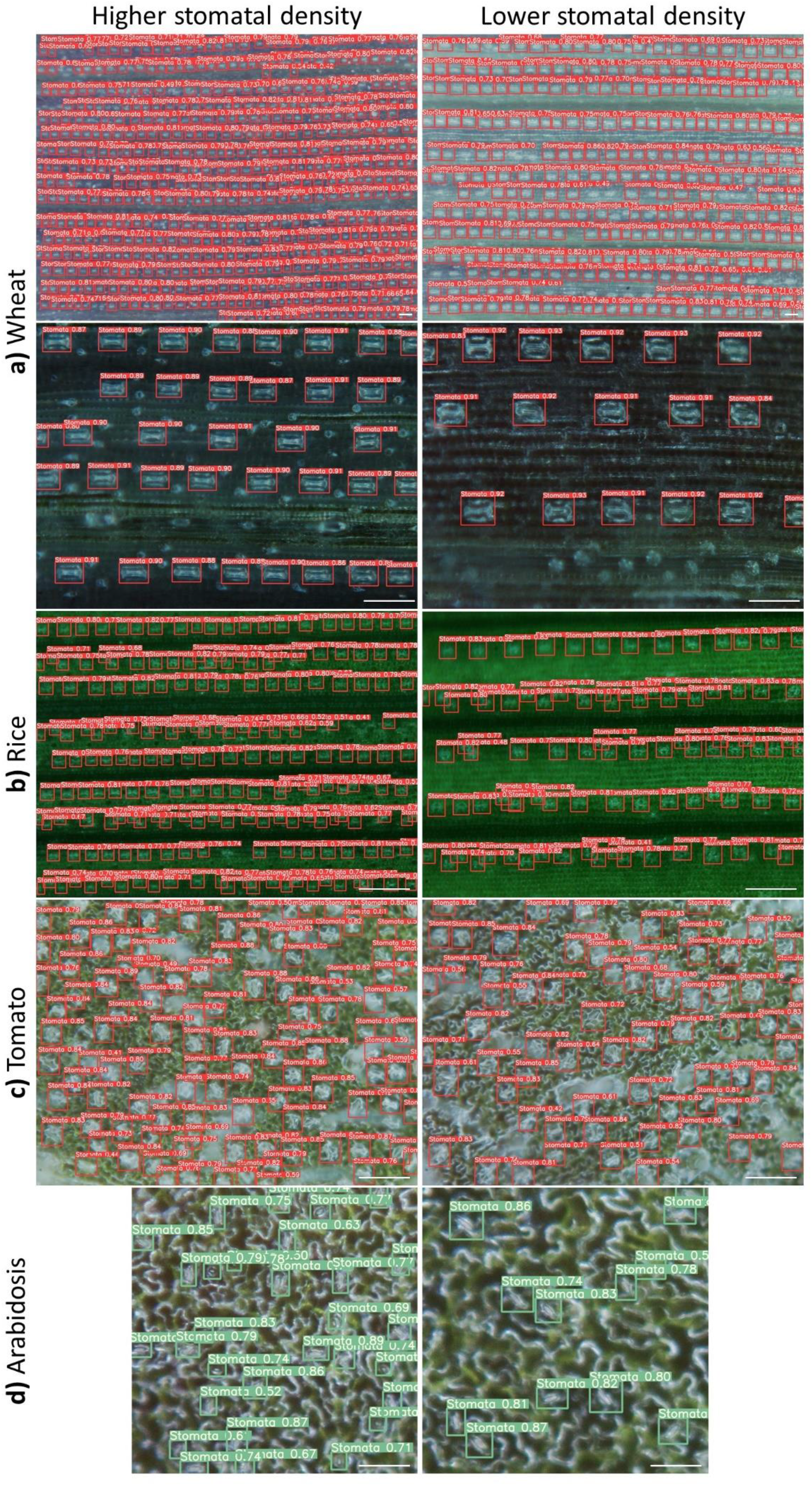
Rapid identification of contrasting stomatal phenotypes: high stomatal density (left) and low stomatal density (right) in wheat (a), rice (b), tomato (c) and Arabidopsis (d). Scale bar = 100 μm.

## Discussion

Research on crop adaptation to biotic and abiotic stresses involves phenotyping large populations to identify germplasm with desirable traits. This requires non-destructive, high-throughput phenotyping tools that accurately measure traits of interest (29). The current low throughput methodology for studying stomatal physiology limits its use in forward genetics screens. Given the central role of stomata in plant physiology, a rapid method was needed to capture the diversity in stomatal traits in plants (24). One of the principles of non-destructive phenotyping is to bring the instrument to the plant, rather than the plant to the instrument. Portable handheld microscopes are therefore convenient tools for non-destructive imaging of the leaf surface. The method described here overcomes the tedious image acquisition process by using a HHM to acquire images directly from the leaf of plants growing in natural conditions. The HHM provided clear images of wheat, rice, tomato and Arabidopsis leaf surface in a few seconds, rather than minutes as is the case with the NP method. Images taken with a HHM can be viewed immediately, and alternative images of the same leaf can be taken if image quality is not satisfactory.

In rice, tomato and Arabidopsis, the small size of stomata only allowed the use of 400× lens, the highest magnification currently available. This magnification provided satisfactory images to count and measure stomata in the three species. Wheat has larger stomata that were visible with 100×, 200× and 400× magnifications. The 100× lens offers a wider field of view and therefore provides a more representative view on stomata number. The 400× lens offers higher resolution of stomata and allows accurate quantification of stomatal size and aperture. The high resolution of the 400× lens makes it possible to record videos of the leaf surface in wheat and to observe and quantify the dynamic changes of stomatal aperture in response to environment, a technique successfully tested in wheat (21). The 200× lens offers a satisfactory compromise between field of view and resolution and can be used to phenotype all stomatal traits in wheat using the same image, which further reduces the time of image acquisition.

This method described here combines the advantages of a portable HHM (rapid, non-destructive acquisition of stomata from leaves on plants in their growing environment), with automated accurate analysis of stomatal features, thus making stomata phenotyping significantly faster than conventional methods. The method was successfully applied to identify contrasting stomatal phenotypes, which is the ultimate purpose of a stomata phenotyping tool. The rapid method is affordable and can be readily used since it does not require specific skills in computer science or programming. The stomata image analysis pipeline is publicly available for each species (30). HHM images can be analysed immediately using the machine learning model.

## Conclusion

Our stomatal phenotyping method provides a rapid, non-destructive tool to determine stomata number, size and aperture if applicable. The experimental setup is portable and allows stomata phenotyping at a large scale, in controlled environments and in the field. This screening tool will accelerate research in stomatal biology in the context of increasing biotic and abiotic pressure on crop production. The method is versatile and can be further adapted to more species using the same HHM and image analysis pipeline.

## Declarations

## Ethics approval and consent to participate

Not applicable

## Availability of data and materials

The datasets generated and analysed during the current study are available in github repository “rapidmethodstomata”, https://github.com/rapidmethodstomata/rapidmethodstomata.

## Authors’ contributions

AE and SR conceived the project. AE, SR and PP designed the experiments. PP and AE performed the experiments. PP, ZZ and AE developed the machine learning model for stomata detection and measurement. PP, ZZ and AE analysed the data. PP, ZZ, SR and AE interpreted the data. All authors drafted the manuscript.

## Acknowledgements

The authors would like to thank Gwen Mayo from Adelaide Microscopy for assistance with the training of staff. We thank The Plant Accelerator, Australian Plant Phenomics Facility (APPF), an Australian Research Facility established under the National Collaborative Research Infrastructure Strategy (NCRIS).

## References

1. Willmer C, Fricker M. The distribution of stomata. In: Willmer C, Fricker M, editors. Stomata. Dordrecht: Springer Netherlands; 1996. p. 12-35.

2. Lawson T, Matthews J. Guard cell metabolism and stomatal function. Annual Review of Plant Biology. 2020;71:273-302.

3. Nunes TDG, Zhang D, Raissig MT. Form, development and function of grass stomata. The Plant Journal. 2020;101:780-99.

4. Melotto M, Underwood W, Koczan J, Nomura K, He SY. Plant stomata function in innate immunity against bacterial invasion. Cell. 2006;126:969-80.

5. Bharath P, Gahir S, Raghavendra AS. Abscisic acid-induced stomatal closure: an important component of plant defense against abiotic and biotic stress. Frontiers in Plant Science. 2021;12. doi.org/10.3389/fpls.2021.615114.

6. Lawson T, Blatt MR. Stomatal size, speed, and responsiveness impact on photosynthesis and water use efficiency. Plant Physiology. 2014;164:1556-70.

7. Franks PJ, Farquhar GD. The mechanical diversity of stomata and its significance in gas-exchange control. Plant Physiology. 2007;143:78-87.

8. Liu C, He N, Zhang J, Li Y, Wang Q, Sack L, et al. Variation of stomatal traits from cold temperate to tropical forests and association with water use efficiency. Functional Ecology. 2018;32:20-8.

9. Liu C, Sack L, Li Y, He N. Contrasting adaptation and optimization of stomatal traits across communities at continental scale. Journal of Experimental Botany. 2022; doi.org/10.1093/jxb/erac266.

10. Ceulemans R, Van Praet L, Jiang Xn. Effects of CO2 enrichment, leaf position and clone on stomatal index and epidermal cell density in poplar (Populus). New Phytologist. 1995;131:99-107.

11. Miyazawa S-I, Livingston NJ, Turpin DH. Stomatal development in new leaves is related to the stomatal conductance of mature leaves in poplar (Populus trichocarpa×P. deltoides). Journal of Experimental Botany. 2005;57:373-80.

12. Zhao W-L, Chen Y-J, Brodribb TJ, Cao K-F. Weak co-ordination between vein and stomatal densities in 105 angiosperm tree species along altitudinal gradients in Southwest China. Functional Plant Biology. 2016;43:1126-33.

13. Reeves G, Singh P, Rossberg TA, Sogbohossou EOD, Schranz ME, Hibberd JM. Natural variation within a species for traits underpinning C4 photosynthesis. Plant Physiology. 2018;177:504-12.

14. Xie J, Wang Z, Li Y. Stomatal opening ratio mediates trait coordinating network adaptation to environmental gradients. New Phytologist. 2022;235:907-22.

15. Zhao Y-Y, Lyu MA, Miao F, Chen G, Zhu X-G. The evolution of stomatal traits along the trajectory toward C4 photosynthesis. Plant Physiology. 2022;190:441-58.

16. Millstead L, Jayakody H, Patel H, Kaura V, Petrie PR, Tomasetig F, et al. Accelerating automated stomata analysis through simplified sample collection and imaging techniques. Frontiers in Plant Science. 2020;11. doi.org/10.3389/fpls.2020.580389.

17. Jayakody H, Liu S, Whitty M, Petrie P. Microscope image based fully automated stomata detection and pore measurement method for grapevines. Plant Methods. 2017;13:94.

18. Meeus S, Van den Bulcke J, wyffels F. From leaf to label: A robust automated workflow for stomata detection. Ecology and Evolution. 2020;10:9178-91.

19. Kwong QB, Wong YC, Lee PL, Sahaini MS, Kon YT, Kulaveerasingam H, et al. Automated stomata detection in oil palm with convolutional neural network. Scientific Reports. 2021;11:15210.

20. Sai N, Bockman JP, Chen H, Watson-Haigh N, Xu B, Feng X, et al. SAI: Fast and automated quantification of stomatal parameters on microscope images. bioRxiv. doi.org/10.1101/2022.02.07.479482.

21. Sun Z, Song Y, Li Q, Cai J, Wang X, Zhou Q, et al. An integrated method for tracking and monitoring stomata dynamics from microscope videos. Plant Phenomics. 2021;2021:9835961.

22. Xie J, Fernandes SB, Mayfield-Jones D, Erice G, Choi M, E Lipka A, et al. Optical topometry and machine learning to rapidly phenotype stomatal patterning traits for maize QTL mapping. Plant Physiology. 2021;187:1462-80.

23. Liang X, Xu X, Wang Z, He L, Zhang K, Liang B, et al. StomataScorer: a portable and high-throughput leaf stomata trait scorer combined with deep learning and an improved CV model. Plant Biotechnology Journal. 2022;20:577-91.

24. Ferguson JN, Fernandes SB, Monier B, Miller ND, Allen D, Dmitrieva A, et al. Machine learning-enabled phenotyping for GWAS and TWAS of WUE traits in 869 field-grown sorghum accessions. Plant Physiology. 2021;187:1481-500.

25. Tzutalin. LabelImg. Git code. 2015; https://github.com/tzutalin/labelImg. Accessed 21 October 2022.

26. Roboflow. https://roboflow.com. Accessed 21 October 2022.

27. YOLOv5. https://github.com/ultralytics/yolov5. Accessed 21 October 2022.

28. Wu Y, Kirillov A, Massa F, Lo WY, and Girshick R. Detectron2. 2021. https://github.com/facebookresearch/detectron2. Accessed 21 October 2022.

29. Furbank RT, Tester M. Phenomics – technologies to relieve the phenotyping bottleneck. Trends in Plant Science. 2011;16:635-44.

30. Rapidmethodstomata. https://github.com/rapidmethodstomata/rapidmethodstomata. Accessed 23 October 2022.

