## Supplementary figures and images for "Rapid non-destructive method to phenotype stomatal traits"

### Supp Fig 1

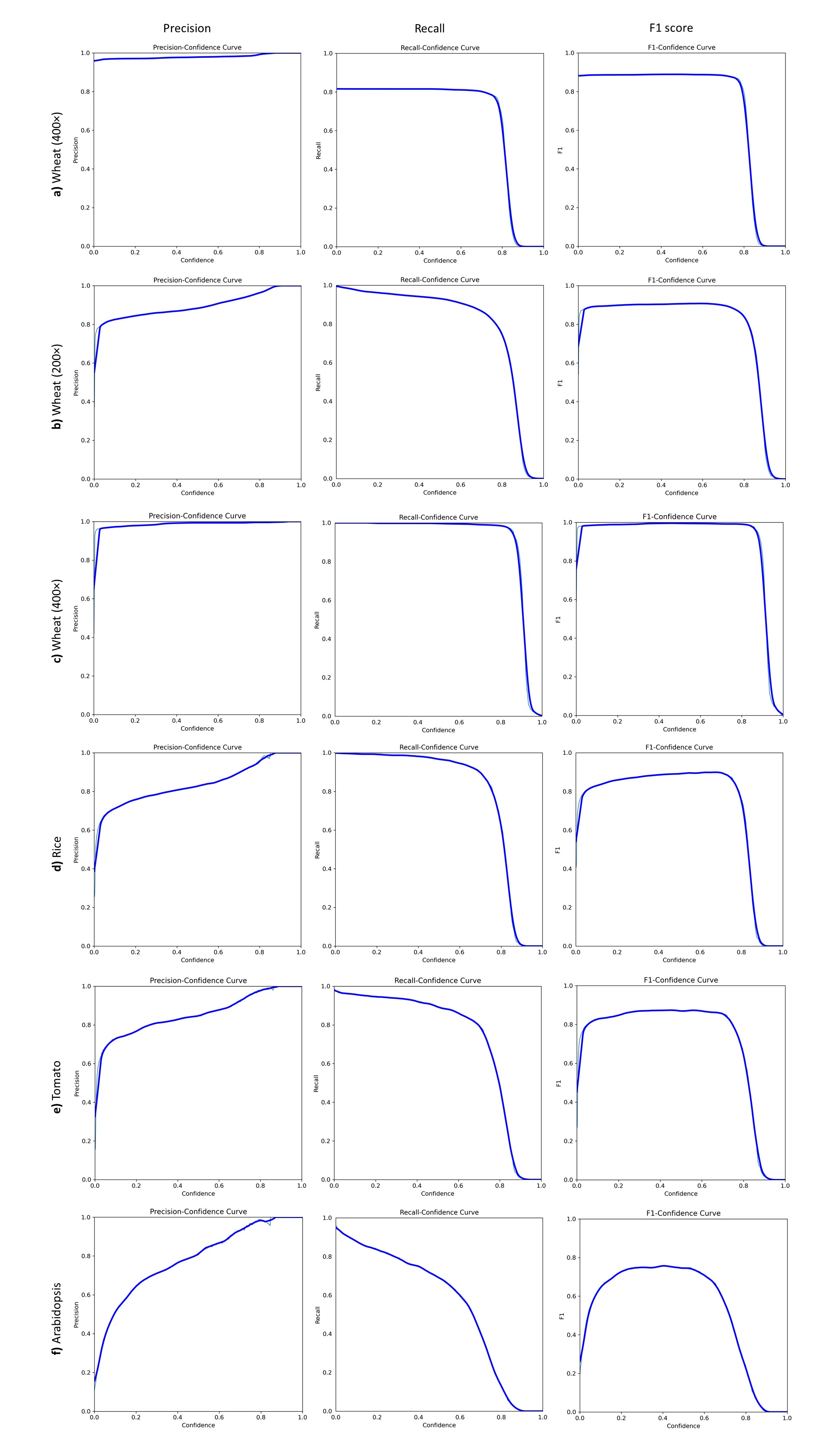
